# Three-dimensional morphodynamics simulations of macropinocytic cups

**DOI:** 10.1101/2020.06.22.165027

**Authors:** Nen Saito, Satoshi Sawai

## Abstract

Macropinocytosis is non-specific uptake of the extracellular fluid playing ubiquitous roles in cell growth, immune-surveillance as well as virus entry. Despite its widespread occurrence, it remains unclear how its initial cup-shaped plasma membrane extensions forms without external physical support as in phagocytosis or curvature inducing proteins as in clathrin-mediated endocytosis. Here, by developing a novel computational framework that describes the coupling between bistable reaction-diffusion processes of active signaling patches and membrane deformation, we demonstrate that protrusive force localized to the edge of the patches can give rise to the self-enclosing cup structure without further assumption of local bending or contraction. Efficient uptake requires an appropriate balance between the patch size and the magnitude of protrusive force relative to the cortical tension. Furthermore, our model exhibits a variety of known morphology dynamics including cyclic cup formation, coexistence and competition between multiple cups and cup splitting indicating that these complex morphologies self-organize through mutually dependent dynamics between the reaction-diffusion process and membrane deformation.

## • Introduction

Macropinocytosis is an evolutionarily conserved actin-dependent endocytic process (1) in which the extracellular fluid is taken up by internalization of micrometer-scale cupshaped membrane ruffles (Fig. 1*A*). A wide range of cell types exhibits macropinocytosis either in a constitutive manner or under growth and other stimulating signals. Macropinocytosis is employed for nutrient uptake in Amoebae *Dictyostelium* (2) and certain cancer cells (3, 4). In immune cells, macropinocytosis plays a role in surveying foreign antigens (5–8). In neurons, macropinocytosis is also employed to regulate neurite outgrowth (9). Understanding the basis of these processes is of biomedical importance due to its link in tumor growth (3, 4), virus entry (10) and spread of prions related to neurodegenerative disease (11). Despite wide occurrence of these phenomena, however, the basic question regarding the very nature of the membrane deformation remains unanswered. The large-scale cup formation involves complex spatiotemporal regulations of signaling molecules and cytoskeletal machineries. Unlike the better-studied clathrin-coated pits, where membrane invagination of ~ 100 nm diameter is formed by clathrin assembly, macropinosome have no apparent coat structures and their size varies between 0.2 – 5*μm* in diameter (6, 12, 13). Furthermore, in contrast to phagocytic cup which extends along the extracellular particles (14, 15), there is no such support to guide the macropinocytic cups externally. These morphological and dynamical features distinct from other endocytic processes indicate a mechanism unique to macropinocytosis that remains to be elucidated.

**Fig. 1:**
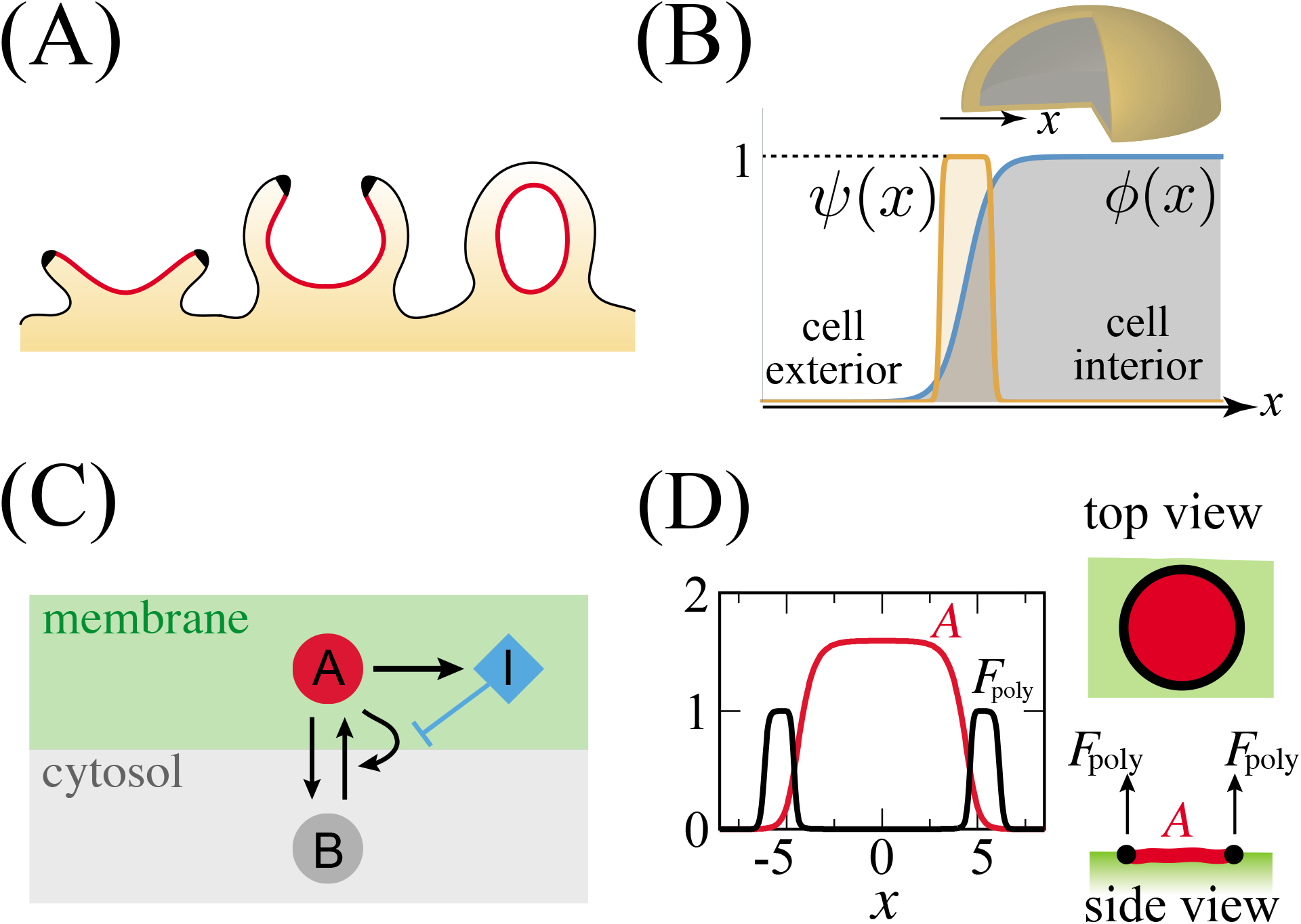
Macropinocytic cup formation and the model schematics. (*A*) Time sequence of macropinocytic cup formation (left to right). A micrometer-scale membrane domain; “active patch” (red) enriched in small GTPases and phosphoinositides grows and expands in the plasma membrane. The Scar/WAVE complex is localized at the edge of a patch (black) (24). (*B*) Phase field *ϕ* defines the state of position *x* in space; occupied (*ϕ* = 1) or vacant (*ϕ* = 0). An auxiliary variable *ψ* is introduced to delineate the border (*ψ* = 1) i.e. the plasma membrane and the rest of the space (*ψ* = 0). (*C*) The schematic diagram of the model reaction. *A* and *B* are active and inactive form of an active patch factor, respectively. ***I*** is a factor that suppresses the positive feedback amplification of ***A*** at the membrane. (*D*) The spatial profile of protruding force *F*_poly_ (Eq.(7)) is determined by the distribution of ***A***. A representative data for a 2D-planar membrane (*K*_0_ = 0.005, *K*_2_ = 0.25 and *n_h_* = 3).

The initial stage of cup formation is identifiable by formation and expansion of an active signaling patch in the plasma membrane that consists of intense accumulation of phosphatydilinositol (3,4,5) tris-phosphate (PIP3) and the active form of small GTPases such as Ras, Rap and Rac surrounded by an edge region enriched in F-actin, Arp2/3 and the Scar/WAVE complex (13, 16). For brevity, we shall refer to this region as ‘active patch’. Relative positioning of these factors remains affixed as the patches grow in size (Fig. *1A* left). The edge of the active patches protrude outward up to several micrometers thus forming the rim of a cup which then curves inward to ingest extracellular fluid (Fig. 1*A* middle). The resulting cup closes by membrane fusion to form a macropinosome (Fig. 1*A* right) which further matures and fuse with lysosomes for degradation of incorporated extracellular solutes (16). The active patch is thought to self-organize by combination of autocatalytic activation of Ras and PIP3 production and their diffusion (17–20).

When observed in the ventral membrane along the substrate, active patches appear as traveling spots and waves – a hallmark of reaction-diffusion mediated pattern formation (17, 18, 21–23). Although these active patches appear to act as a prepattern or ‘template’ for macropinocytic cup (24), little is known how these materialize into the formation of the cup itself.

In recent years, progress in theoretical and computational approaches have allowed one to address dynamical properties of cellular- and sub-cellular scale membrane deformation such as amoeboid motion, filopodia formation. Common to these modeling approaches is mathematical formulation that describes the underlying regulatory kinetics together with a moving boundary. This physico-chemical coupling makes the problem unique and challenging, since the very nature of highly deformable boundary requires elaborate techniques to solve the interface physics that are often computationally laborious and expensive. Many of the studies have focused on cases that can be approximated in one- and two-dimensional space including but not limited to formation of filopodia during axonal elongation (25), pseudopodium in ameboid migration (26), lamellipodia of fish keratocytes (27–29), while relatively few attempts have been made for 3 dimensional dynamics (30–32). Models of 2-D dynamics by the active patches constrained to the ventral (18) or the dorsal-side (19) of the plasma membrane has been analyzed. Given its geometry, understanding the full nature of membrane deformation in macropinocytosis poses a challenge that requires full 3-dimensinal modeling with topological changes in membrane. In this paper, we propose and analyze a minimalistic 3-D model to address the relationship between the self-organizing active patches and the geometry of macropinocytic cup formation and closure. Our results indicate that relative simple rule of self-organization coupled with membrane protrusion can explain the entire sequence of the dynamics starting from patch expansion, cup formation to cup closure without further need for specialized machineries to regulate local curvature.

### Model

We adopt a modeling strategy that combines two elementary processes: (i) deformation of the membrane and (ii) reaction-diffusion process of signaling molecules on the deformable membrane. To describe the membrane, we employ the phase-field method, which allows one to simulate interfaces with complex geometry such as growing crystals (33, 34), vesicle coarsening or fission (35) as well as an overall shape of migratory cells (18, 20, 25, 27). The phase-field approach allows one to compute cellular membrane deformation on the order of micrometers in spatial scale and seconds to minutes in timescales, which is in contrast to nanometer-scale models that describe microsecond order phenomena (36). Here, an abstract field variable *ϕ* is introduced to describe the cell interior region *ϕ* =1 and the exterior region and *ϕ* = 0 (Fig. 1*B*). *ϕ* is assumed to be continuous and varies sharply at the interface with finite width characterized by a small parameter *ε*. Following previous studies (18, 27), here, we adopt the following equations (see SI Text for derivation)

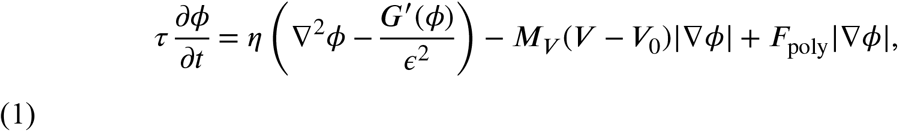

where *G*′ = 16*ϕ*(1 – *ϕ*)(1 – 2*ϕ*) and *V* = ∫ *ϕd**r***. The first term in the right hand side represents curvature-driven force associated with surface tension *η*. The second term imposes a constraint on the cell volume to 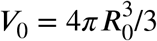 where *R*_0_ is the cell radius and *M_v_* is a constraint parameter. The third term describes the force normal to the interface driven by dendritic actin polymerization. The magnitude of force *F*_poly_ is assumed to be a function of the local concentrations of signaling molecules as described below.

For time development of the signaling molecule, let us assume an interconversion between the active form ***A*** on the plasma membrane and inactive cytosolic form 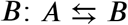 (Fig. 1*C*). The total number of molecules is fixed to ***A***_t_, and the conversion from ***A*** to ***B*** is assumed to take place at a constant rate, whereas that of ***B*** to ***A*** is facilitated in an autocatalytic manner. This scheme gives rise to bi-stability, where ***A*** takes two states: zero and a finite positive value. When ***A*** is locally perturbed from ***A*** =0, a domain where increase in ***A*** takes place spreads in space and eventually stops due to depletion of ***B***, thereby creating a stable spot pattern which we shall consider as a mathematical representation of an active patch. Let us further introduce a factor ***I*** that inhibits conversion of ***B*** to ***A*** so that the active patch has a finite lifetime. The above basic reactions are expressed in the following dimensionless reaction-diffusion equations (see SI Text for derivation):

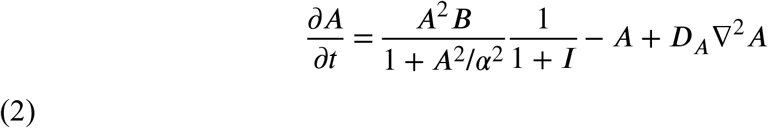

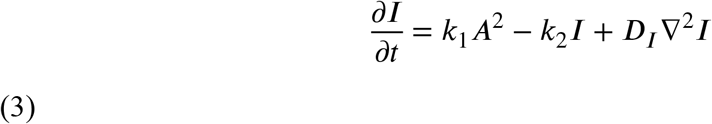

where ***D_A_*** and ***D_I_*** are diffusion constants of ***A*** and ***I*** molecules, respectively. The parameter *α* dictates a half saturation concentration of the Hill function in the autocatalytic reaction ***B*** → ***A***. The second equation assumes a negative feedback that produces the inhibitor *I* at the rate *k*_1_***A*** and degraded at a constant rate ***k***_2_. When diffusion of ***B*** molecule is sufficiently fast, ***B*** = ***A***_t_/*S* – 〈***A***〉, where ***S*** is the cell surface area ***S*** = ∫ *ψ/ε* dr^3^ and 〈***A***〉 is the total of ***A*** divided by ***S***. Note that ‘*A*’ and ‘*B*’ can also be membrane-bound factors as long as diffusion of ‘*B*’ is sufficiently fast compared to that of ‘*A*’. For ***I*** = 0 and *k*_1_ = 0, the reaction-diffusion equations are reduced to the well-studied wave pinning model of cell polarization (37, 38).

To define spatial coordinates occupied by the plasma membrane, let us introduce an auxiliary phase-field 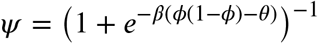 which specifies the interface between cell exterior (*ϕ* = 0) and interior (*ϕ* = 1) region. *ψ* takes constant value *ψ* =1 at the cell membrane and *ψ* = 0 elsewhere (Fig. 1*B*). Here, *β* takes a sufficiently large value so as to render the interface between inside and outside of the membrane sharp. *θ* is set so that the *ψ* is non-zero at the interface of *ϕ* with thickness *ϵ*. The unique aspect of the present approach is the introduction of this auxiliary field *ψ* thereby allowing Eqs. (2) and (3)) to be solved numerically at the interface only. In contrast, previous 2D models (18, 27, 28) made distinction only between occupied (cell; *ϕ* =1) and vacant (no-cell; *ϕ* = 0) regions and assumed reaction that take place throughout the occupied space. Using *ψ*, we arrive at the following equations:

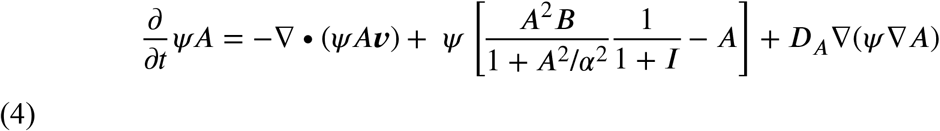

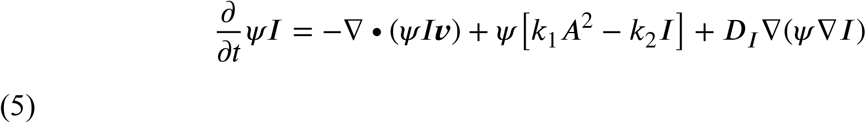

where the first terms in the right hand side are the advection term and ***v*** is given by

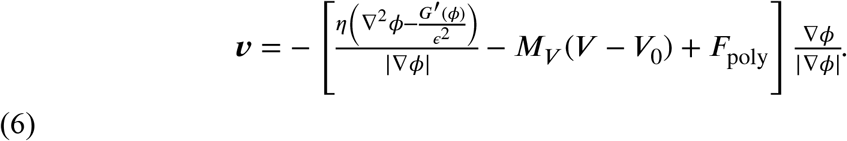

Given that protrusive actin filaments are concentrated at the edge of the activated patch (16, 24), we assume that protrusion is facilitated when *A* is within a certain range as illustrated in Fig. 1*D*. To implement this, the actin-dependent force generation in Eq. (1) and (6) are given as in the form of the force term

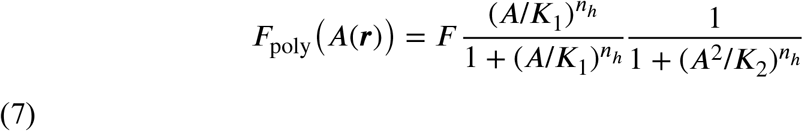

so that *F*_poly_(*A*) ~ *F* for 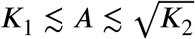.

## • Results

### Mutually dependency between the patch dynamics and deformation drives cup formation and closure

First, we demonstrate an overall time development of the 3-dimensional model in the absence of the inhibitor by setting *k*_1_ = 0. As an initial condition, we chose a membrane sphere with *A = I* = 0 except for a small circular region with radius *r*_init_ where the local concentration of *A* takes random value from 0 to 5.0 on each grid. Representative results are shown in Fig. 2*A* (see also Movie S1). Due to bistability, a local active patch defined by high *A* begin to invade the basal state of low *A* as a propagating front (Fig. 2*A* *t* = 4 orange region). As the patch expanded, the membrane protruded at the patch periphery and formed a cup shaped circular extension (Fig. 2*A*; *t* = 4 – 24 orange and green border). After the patch grew to a certain size, the expansion slowed down. At the same time, the protrusion formed an overhang while the center of the patch curved slightly inward to form a cup (Fig. 2*A*; *t* = 24 – 44). The rim of the cup shrunk and annihilated as the membrane sealed itself to completely surround a large volume of extracellular space (Fig. 2*A*; *t* = 65). The coordinated manner in which a circular ruffle encircling a non-protruding area extended, shrunk and closed showed a close parallel to the cup dynamics observed in *Dictyostelium* (24). Also of note is the marked accumulation of *A* in the inner territory and its exclusion from the rim which are in good agreement with the patterns of bona fide active patch marker PIP_3_ and Ras-GTP (24, 39–41).

**Fig. 2.**
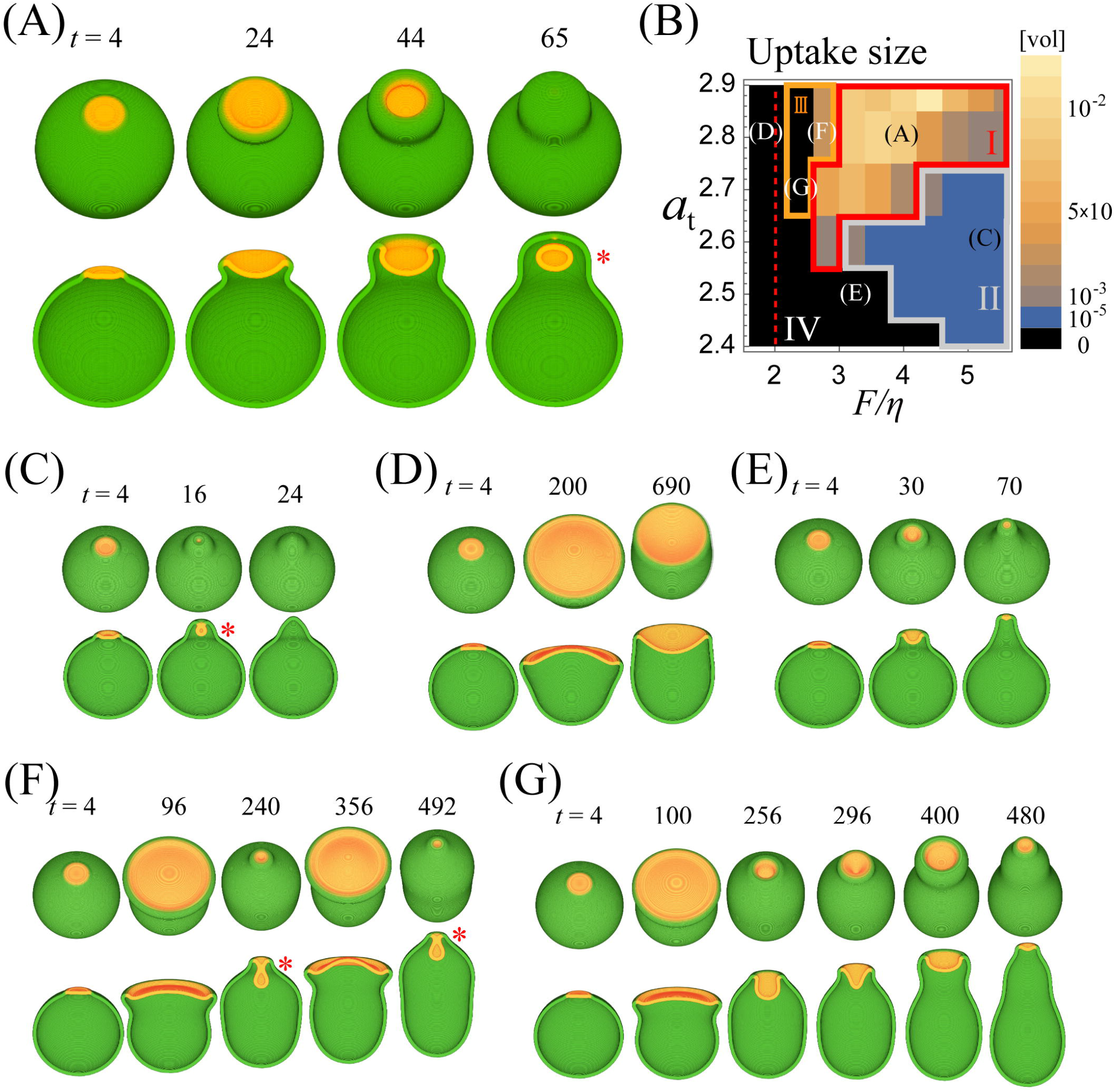
Membrane protrusion at the edge of an active patch is sufficient for the formation of the basic cup-like structure and its closure. Simulation results: (*A*) a representative time course of the numerical simulations (*F/η* = 4.0, *a*_t_ = 2.8). The active patch (red; ***Aψ*** > 0) and the membrane (green; *ψ* > 0) shown as merged RGB images; birds-eye view (upper panels) and as cross sections along the median plane (lower panels). Asterisks indicate cup closure. (*B*) Phase diagram of the cup dynamics. Color bars indicate the volume of enclosure normalized by the cell size (blue to yellow). The cutoff volume for successful cup closure was set to < 10^−5^ (black). Averages of six independent simulation runs (three of each for ***r***_init_ = 1.0 μm and 1.5 μm) are shown. Phase I and II: enclosure in all or part of the six trials, respectively. Phase III: repetitive cup formation. Phase IV: cup closure failed in all simulations runs. The red dashed line is the estimated minimal force ***F/η*** = 2.0 required for protrusion. Parameter sets in (*A*) and (*C-G*) are indicated in the diagram. (*C-G*) Representative time course for (*C*) ***F/η*** = 5.2, *a*_t_ = 2.6 (*D*) ***F/η*** = 1.6, *a*_t_ = 2.8, (*E*) ***F/η*** = 3.2, *a*_t_ = 2.5, (*F*) ***F/η*** = 2.8, *a*_t_ = 2.8, (*G*) ***F/η*** = 2.4, *a*_t_ = 2.7. Other parameters: ***τ*** = 10, *D_a_* = 0.1, *α* = 1.0, *ε* = 0.8, ***M_V_*** = 5.0, *β* = 100.0, *θ* = 0.105, *η* = 0.5.

Whether the cup closed or not depended on the parameters and the initial condition. Cup closure was judged by evaluating whether the region with *ϕ* = 0 surrounded by *ϕ* =1 based on quasi 3-dimensional simulations where dimensionality was reduced in an axis-symmetric coordinate for easier detection of morphology criteria and computation involving exhaustive parameter search (fig. S1 *A-H*; see also Methods). In the phase field framework, the topological change that accompanies membrane fusion is naturally established by simply solving the partial differential equation Eq. (1) without additional numerical implementation. For enclosure of a large volume, the critical parameters were the ratio between force per unit area *F* and the surface tension *η* i.e. *F/η*, and the total amount of ***A*** and ***B*** per unit area 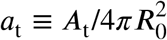. Figure 2*B* illustrates a phase-diagram for cup closure. Black regions represent parameters that were unable to support enclosure, otherwise color represents the enclosed volume relative to the cell volume *V*_0_ (Fig. 2*B*). A similar portrayal of the parameter space was obtained based on the elapsed time between the patch initiation and the cup closure (fig. S2*A*) and the ingestion efficiency (fig. S2*B*). All phase diagrams were obtained by averaging results from two initial patch size *r*_init_ (fig. S2*C, D*). The parameter space could be divided into four domains: Phase I – IV based on the success rate of closure. Phase I includes the example shown in Figure 1 *A* where parameters supported enclosure in all cases. Phase II consists of parameters where cup closure depended on the initial conditions. Here, due to small patch size, the cup and hence the enclosed extracellular volume was sometimes extremely small (Fig. 2*C*). In Phase IV, cup closure failed for all simulations runs (Fig. 2*D, E*).

Requirements for successful cup formation and closure (Phase I) can be understood from the characteristic dynamics observed when cups failed to support large volume uptake. Phase IV consisted of two patterns of incomplete closure depending on the value of *F/η*. When *F/η* was small, patches and cups persisted indefinitely without shrinking or closing (Fig. 2*D*) for both high and low *a*_t_. At high *F/η*, cup shrunk without closing when *a*_t_ was not high (Fig. 2*E*). The behavior at low *F/η* was due to lack of sufficient protrusive force for cup development. Consider a cross section of a protrusion with width *2R* (fig. S3), the force per unit length *F* exerted on the semicircular head of length *l = π****R*** should be twice as large as the line tension *η* required to maintain the protrusion. Hence, the minimal force *F*\* must obey *F*\*/*η* = 1/***R*** = *π/l*. Based on cortical tension of ~ 0.7 nN/μm (42) and an estimate for protrusive force ~ 6.5 to 9 nN/μm^2^ (protrusive force by a single microfilament 5 to 7 pN times the filament density (43)), the condition *F/η* > 1/***R*** is satisfied for protrusion width of ≿ 0.2 μm. While *l* in real cells has not been measured quantitatively, projections thinner than 0.2 μm would require larger ***F/η*** than the above estimate. Relative ease of imaging the cups under the conventional confocal microscope suggests they are above the diffraction limit (> 0.25 μm) which is within this force requirement. Because the spatial resolution of our numerical simulations were limited by the computational time, for systematic parameter studies, parameters ***K***_1_ and ***K***_2_ in Eq. (7) were chosen so that ***l*** ~ 1.5 *μm* (Fig. 1*D*, black plateau; fig. S3), hence ***F***\*/***η*** ~ 2.0 *μm*^−1^ which is consistent with the boundary in the phase diagram (Fig. 2*B*; red dashed line). In contrast to the force constraints at small ***F/η***, the characteristic behavior at high ***F/η*** (Fig. 2*E*) was due to lack of sufficient patch size at small *a*_t_. Here, the resulting small cups gave rise to high negative curvature which in turn provides strong restoring force in the inner territory that prevented the protrusion from curling inward. This resulted in a shmoo-like cell morphology (Fig. 2*E*, *t* = 70) which eventually returned to symmetric sphere as the patch disappeared. This patch attenuation was a distinct feature that arose due to self-consistency requirement that the edge of the patch must define the point of protrusion and vice-versa. If protrusion were to come close and coalesce due to high tension, the region that it surrounded must also disappear. One should note that the same parameters support a persistent patch if it were not for deformation, thus the coupling of reaction-diffusion process and deformation is essential.

In Phase III, cup formation was observed to repeat at the same site. A similar behavior has been observed in the standard axenic strain of *Dictyostelium discoideum* and in an even more pronounced form in the null-mutant of RapGEF *(gflB)* (44). In our simulations, there were two patterns of repetition both of which occurred under conditions that allowed formation of exceedingly large cup (Figs. 2*F* and *G*). In the first examples (Fig. 2*F*; see also Movie S2), cup closed at its waist (Fig. 2*F*; *t* = 240 sec) while the remaining open half continued to expand at the edge (Fig. 2*F*; *t* = 356). After the second closure (Fig. 2*F*; *t* = 492), the rim disappeared and there was no more cup formation. The other pattern occurred for slightly weaker force (Fig. 2*G*; see also Movie S3). Here, cup closure was stalled in the middle (Fig. 2*G*; *t* = 296) as the patch continued to expand laterally before the next attempt at the cup-formation (Fig. 2*G*; *t* = 400). While distinction between these two behaviors is difficult to resolve experimentally, the markedly elongated cell shape (Fig. 2*F*; *t* = 492 and Fig. 2*G*; *t* = 480), and the lengthening of time required for enclosure (fig. S2A) are in accordance with what has been reported for the *gflB* mutant. The size of the Phase III region depended on the time scale of deformation ***τ***. In the examples shown above (*τ* = 10 sec), normal cup closure (Phase I) was predominantly observed, and Phase III was confined to a narrow domain between Phase I and IV (Fig. 2*B*). For smaller *τ* (*τ* = 5 sec), Phase II became dominant, and Phase I and III were both confined to narrow regions in the parameter space (fig. S4*A*). In contrast, at large *τ* (= 20 sec), the Phase I and III regions expanded (fig. S4*B*). Overall, normal cup closure (Phase I) is realizable at large τ, however it comes at the cost of also inviting repetitive dynamics that are often incomplete (Phase III) in addition to the overall process slowing down (fig. S4*B* middle) making the process less efficient (fig. S4*B* right).

### Inhibitor and mass conservation determines duration of the patch and cup dynamics

Large cell-size cups are frequently observed in the axenic strain of *Dictyostelium* (24, 40, 45), however they do not exist indefinitely. The active signaling patches are mostly transient and eventually vanishes with a lifetime of few minutes (17, 18, 22, 23, 46). In our simulations, the active patches on their own have finite lifetime when the presence of the inhibitor ***I*** is non-negligible (*k*_1_ ≠ 0). For *k*_1_ = *k*_2_ = 2.0 × 10^−4^, the inhibitor ***I*** increases at a much slower timescale than the initial expansion of the active patch. Eventually, ***I*** becomes high enough to suppress *A* i.e. the activate patch (fig. S5) when *a*_t_ satisfies a certain condition (see SI Text). In Phase III, the presence of the inhibitor repressed the repetitive cup formation and abolished the ruffle formation (fig. S5*C* and *F*), whereas no change was observed for Phase I and II.

Besides the inhibitor, the assumed mass conservation of the signaling molecule can also prevent futile formation of excessively large cup. This effect becomes most evident when there are simultaneous and constitutive occurrence of active patches. Let us examine slightly complex situations where activation of *A* is allowed to occur at random positions ***x***_c_ at rate θ per volume. The spatial profile of noise follows 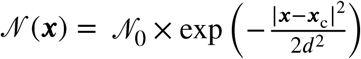, where ***d*** is the initial nucleation size, and 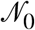 is the noise intensity that follows an exponential distribution with the average *σ*. Figure 3*A* shows representative snapshots from independent simulation runs (see also Movie S4). A new active patch was nucleated before existing cups closed thus allowing multiple cups to coexist. Depending on the size and amplitude of the noise, some cups closed successfully, while others shrunk and vanished before they can close. Incomplete closure occurred even when the same parameter supported closure for an isolated cup (Phase I). This can be explained by effective lowering of *a*_t_ available per cup. Due to continual cup formation and closure, the cell shape deviated markedly from the initial sphere and took complex and processive morphology that highly resembled axenic strains of *Dictyostelium*. In the parameter regime that supported relative large and slow cup closure (Phase III), these features became more exaggerated (Fig. 3*B*). Multiple cups were indeed frequently observed in *Dictyostelim* cells, and they either successfully closed to form endosomes or vanished without closing (40).

**Fig. 3:**
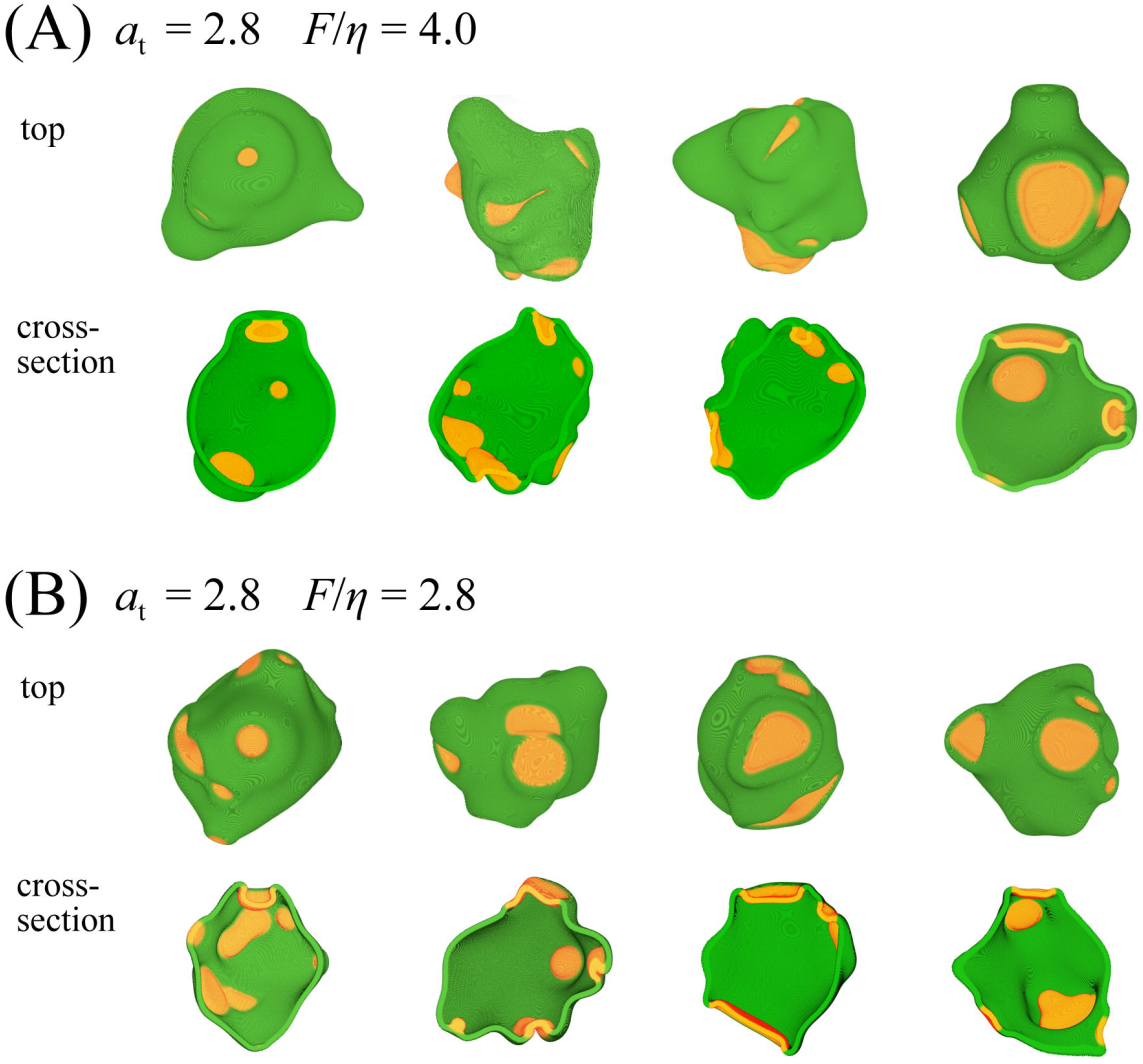
Complex cell morphologies result from multiple stochastic patch initiation. (*A, B*) Representative snapshots from independent simulations with the same parameter sets as Fig. 2*A* and *F* (*I* = 0). Top overhead view (upper panels) and the midline cross section (lower panels) with merged RGB images (Green: cell membrane (*ψ* > 0). Red: active patches (*Aψ* > 0). Noise parameters: *σ* = 8.0, *d* = 15.0, *λ* = 3 × 10^−5^.

### Excitability arises in the presence of strong inhibitory signal and drives cup splitting dynamics

The cup dynamics described above was monotonous, meaning that the initial active patch more or less dictated when and where a cup formed, and it grew due to bistability until it consumed all ***B***. In *Dictyostelium,* however, cups are known to also multiply or reduce in number by splitting and coalescence of existing cups (24). In the present model, when the production of ***A*** is no longer a saturating function (large *α* in Eq. 2), the active patch (a region with high *A*) can become out of phase with a high ***I*** region. As a consequence, the region occupied by high ***I*** will trail behind a moving active patch and can disrupt it (fig. S6*A*, *B*). To study this behavior in detail, let us consider a case *α* → ∞ so that Eq.(2) now becomes

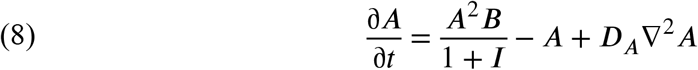

which is the same equation introduced earlier as a part of a model for the patch dynamics in circular dorsal ruffle (19). The key difference in the present model, apart from incorporation of the membrane deformation, is that Eq. (8) is coupled to Eq. (3) with quadratic dependence on *A* which is essential for providing a rich behavior as follows. The equation has three different parameter regimes: mono-stable, bi-stable, and excitable (Fig. 4*A*, see also fig. S7*A-C* for finite *α*). In the excitable regime, null-cline analysis (Fig. 4*B*, left panel) shows that, for a small 〈***A***〉 (i.e., for 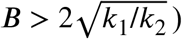), small perturbation from the fixed point ***A*** = 0 gives rise to a large excitation of *A*. For large 〈***A***〉 (i.e., for 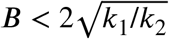: right panel in Fig. 4*A*), excitability disappears and ***A*** falls immediately to the basal state even when strongly perturbed. Interestingly, this in turn brings the system back to an excitable state hence ***A*** is again easily perturbed and brought transiently to a high level. In other words, depending on 〈***A***〉 i.e. the total size of active patches, excitability is switched on and off in a sequential manner. This switching of excitability destabilizes the expanding front of active patches (fig. S6*A*), similar to splitting patches or waves observed in the ventral side of the plasma membrane (17, 18, 21–23).

**Fig. 4:**
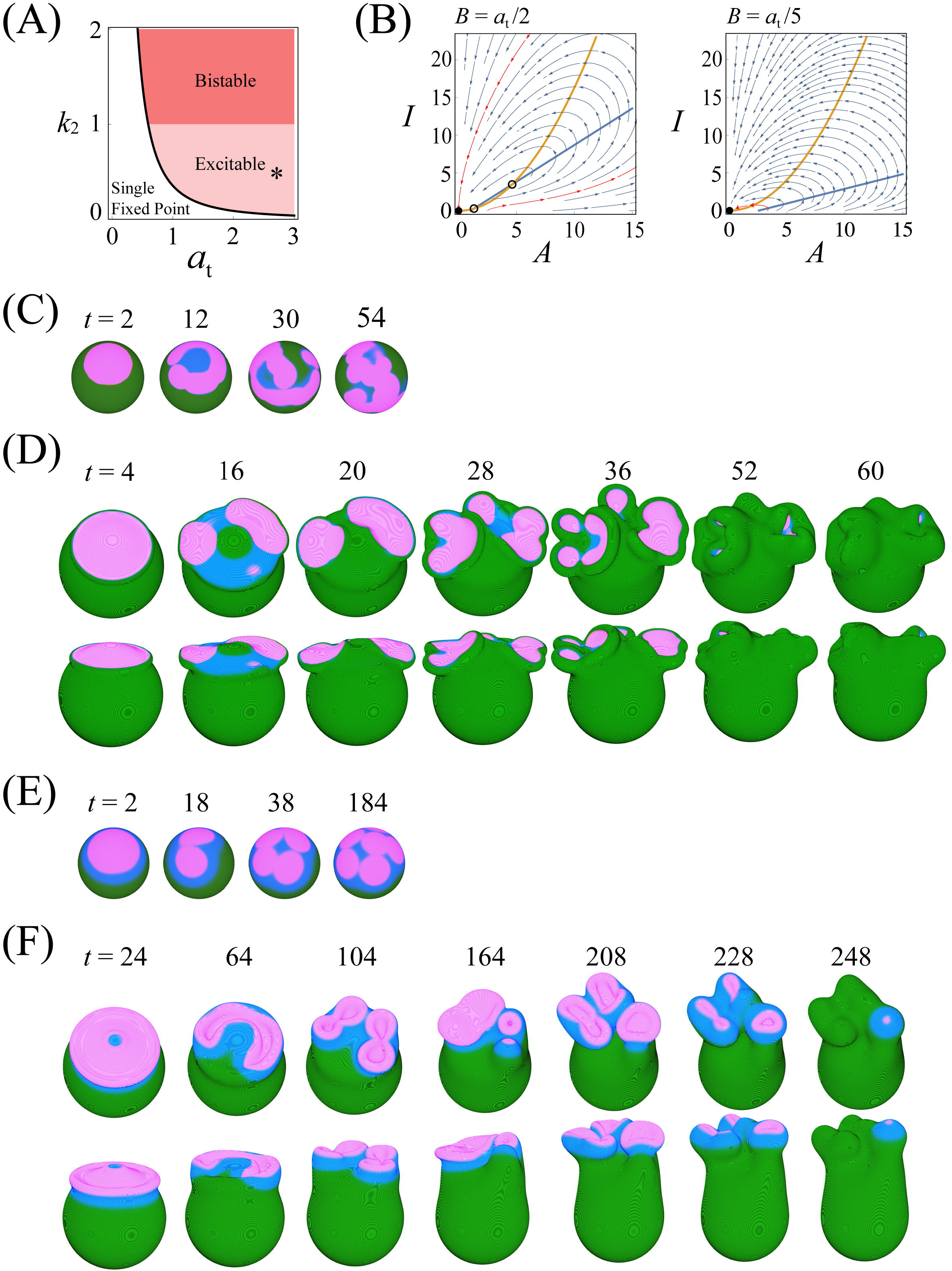
Presence of an inhibitor gives rise to cup splitting dynamics. (*A*) Phase diagram of chemical reaction Eq.(2) decoupled from deformation dynamics in the presence of inhibitor kinetics (Eq.(3)) for *k*_1_ = 0.088. Depending on *k*_2_, the system is bi-stable (*k*_2_ > 1) (red region) or excitable (*k*_2_ < 1) (pink region). A single fixed point ***A*** = 0 for 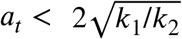 (white region) and three fixed points ***A*** = 0, and 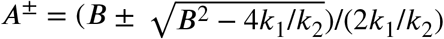 for 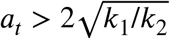 (pink and red regions). ***A*** = 0; stable. ***A^−^***; unstable. ***A^+^***; stable (*k*_2_ > 1) or unstable (*k*_2_ < 1). (*B*) Null-clines in the excitable regime for 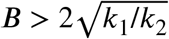 (left panel) and 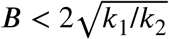 (right panel). Fixed points (Filled circle: stable. Open circle: unstable). Excitatory trajectories (red arrows) invoked by small perturbation to ***A*** = 0. (*C, D*) Representative dynamics (*a*_t_ = 1.985, *k*_1_ = 0.088, *k*_2_ = 0.54, *D_a_* = 0.085 and *D_i_* = 0.11) on a fixed spherical field (*C*) and deforming membrane (*D*) (*τ* = 7.0, ***F*** = 3.7, ***K***_1_ = 0.01, ***K***_2_ = 0.1 and *n_h_* = 5). (*E, F*) Representative dynamics (*a_t_* = 1.94, *k*_1_ = 0.088, *k*_2_ = 0.54, *D_a_* = 0.26 and ***D_i_*** = 0.87) on a fixed spherical field (*E*) and deforming membrane (*F*) (***τ*** = 20.0, ***F*** = 3.0, ***K***_1_ = 0.086, ***K***_2_ = 1.8 and *n_h_* = 3). Other parameters are same as in Fig. 2.

When coupled to membrane deformation, however, a broad protrusive force profile in the region surround by a patch (fig. S6*C* and *D*; *t* = 8, 0 < *r* < 3) smoothed out the fragmented active patches before daughter cups developed (fig. S6*E* and *F*). While this can be circumvented at small *F,* fragmented cups then failed to close due to lack of sufficient protrusion (fig. S6*G*). A recent CryoEM study of the ventral actin waves demonstrated that the form and alignment of actin filaments at the edge of the patch and those in the inner region are distinct and thus hints at the presence of debranching factors that trails behind the expanding edge (47). Such notion is line with sharp localization of Scar/WAVE complex at the edge of a patch (24) and depolymerization factor Coronin at the rear of the edge (Bretschneider et al 2009. Biophys J). To study such an effect in the model, let us modify the force term so that that *I* not only suppresses amplification of *A* but also competitively inhibits force generation by *A*, so that

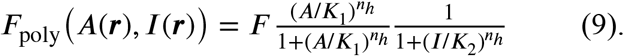

Note that the original form of *F*_poly_ (Eq. 7) is recovered when ***I*** is at the steady state; i.e. ***İ*** = 0, and ***D_I_*** is negligible. The periphery of the active patch is defined by high ***A*** and low ***I***, thus under Eq. (9), the force profile became restricted to the edge (fig. S6*H, I; t* = 8). Protruding force in the inner territory only appeared later to surround the split patches (fig. S6*H*; *t* = 15). Accordingly, expanding active patches broke up repeatedly while some of the daughter patches quickly merged with existing ones and gave rise to cup-shaped circular ruffles (Fig. 4*D*; *t* = 20, 28, and Movie S5). By ruffles, we mean that the rim of cup was no longer smooth and circular but more undulated and complex in shape. Splitting of an activate patch during ruffle formation causes its fragmentation (Fig. 4*D*; *t* = 36). A notable difference from multiple cups occurring in non-excitable regime (Fig. 3) was that multiple patches and cups continued to emerge starting from a single founder. These sequence of events and their appearance; splitting followed by formation of cup-shaped ruffles (Fig. 2*D*) are remarkably similar to how, in *Dictyostelium,* an active Rac and F-actin rich region expands together with membrane ruffles then become fragmented into multiple macropinocytic cups (24).

## Discussion

The present work suggests an unexpectedly simple yet concerted mechanism that underlies formation and closure of macropinocytic cups. First, a locally activated signaling patch represented by high ***A*** in the model appears. The active patch expands in a self-organized manner via autocatalytic transition of bistable nature from the state of low *A* to high *A*. From there, two key assumptions in the model dictate the fate of the patch and the resulting cup. (i) Growth of an activated patch is limited due to the finite amount of the signaling molecules (Eq. (2)); i.e. the sum of ***A*** and ***B*** molecules ***A_t_*** (or *a*_t_ in the normalized form) is fixed within a cell. (ii) Protruding force is restricted to the edge of an active patch (Eq. (7) and (9)) (16, 48). Due to the constraint (i), a patch first expands (fig. S8; *t* = *t*1), then slows down as it reaches its size limit (fig. S8; *t* = *t*_2_). The edge continues to protrude and forces the patch area to expand. However because *B* is no longer available, *A* at the patch boundary must be brought down to the low state. Thus the position of the patch boundary (fig. S8; *t* = *t*_3_, black circle) is effectively displaced from the rim of a cup (fig. S8; *t* = *t*_3_, blue asterisk) towards the inner territory (fig. S8; *t* = *t*_3_). Because protrusive force is generated at the patch boundary (ii), the protrusion begins to curve inward, forming an overhang (fig. S8; *t* = *t*_4_, *t*_5_) and continues to advance until they meet each other. We should note that spontaneous curvature is assumed to be negligible in the present formulation (Eq. (1)), and that the involution arises due to mutuality between the reaction-diffusion process and deformation dynamics in defining the position of the protrusion. In this light, the work brings to light a distinct mechanism of membrane invagination that contrasts with those driven by local curvature; e.g. formation of endocytic vesicles by clathrin (49) and BAR-domain containing proteins (50, 51).

The high similarity between the range of complex morphology dynamics observed in the present simulations and those in *Dictyostelium* cells suggest that the kinetics adopted in the current model captured the essence of the underlying regulation and the cell mechanics. The critical parameter that determined the occurrence of a patch and its size was *a*_t_. Strong candidates for ***A*** and ***B*** are active and inactive form of small GTPase such as Ras, Rap and Rac or their upstream and downstream signaling partners such as PI3K which are all found enriched in the activated patch (24, 39, 52). PI3kinase requires Ras binding for its activity (53, 54) and thus the variable ***A*** may represent Ras in complex with PI3kinase or its product PIP3 and the variable ***B*** may be regarded as their inactive forms. In fibroblasts, microinjection of active Ras protein induces macropinocytosis (55). RasS mutation in *Dictyostelium* cells are known to inhibit macropinocytosis (39, 56). These perturbations can be understood from increasing or decreasing *a*_t_ and hence the size of the active patch. Due to non-dimensionalization in Eq. (2), lowering of *a*_t_ can also result from decrease in the autocatalyic reaction ***B → A*** (see SI Text). The analysis is in line with a recent suggestion based on the observation of smaller patches and macropinosomes in PI3kinase mutants (24, 57) and in a double mutant of Akt/PkbA and PkbR1 (57) that there likely is a positive feedback loop between PIP3 production and its downstream PKB in *Dictyostelium* (57).

Apart from bi-stability, our model suggests that excitability arises when the self-amplification of ***A*** is less saturated (i.e., large *α*). In *Dictyostelium,* loss of Ras GTPase-activating protein (RasGAP) Neurofibromin (NF1) causes formation of oversized macropinosomes, increases fluid uptake and facilitates cell growth in liquid media (45). In our model, a decrease in RasGAP would correspond to lowering the rate of reaction ***A → B***. Due to parameter non-dimensionalization in Eq. (2), not only *α*_t_ but also *α* increases in this case (see SI Text). Since elevation in *α* brings the system to the excitable regime, attenuation of RasGAP makes it an ideal point of perturbation to enhance macropinocytosis; i.e. an increase in the number of patches due to splitting in addition to supporting a larger patch size. Both expression of activated Ras in the wild-type cell (41) and Ras-GAP mutation (45) are known to enhance fluid uptake. Our model assumes that the inhibitor ***I*** weakens autoregulatory amplification of ***A***. From ***İ*** = 0 at Eq. (3), one can see that ***I*** imposes saturation in the production of ***A*** even at high *α*, and thus has a similar effect to changing *α*. In addition, ***I*** acts critically for patch duration as well as for cup splitting (Fig. 4*D*). For large *a*_t_, absence of the inhibitor ***I*** caused cup formation to repeat at the same site due to incomplete closure of an oversized cup (Fig. 2*B*; Phase III). Following the line of thoughts that ***A*** maybe regarded as an activated form of small GTPase, ***I*** would be a factor that suppresses guanine nucleotide exchange factor (GEF)s. In line with the model behavior, a knockout of Ras/Rap GEF (GflB) does indeed repeat cup formation at the same site (44).

For efficient uptake, the key mechanical parameter was the magnitude of protrusive force relative to that of the cortical tension. Our model predicts that high *F/η* should facilitate macropinocytosis, which is in line with increase in macropinocytosis under decreased membrane tension (58). Although the variable ***A*** and ***I*** are abstract and collective representation of regulatory factors, from the cell mechanics point of view, they must be closely linked to the nucleator Arp2/3 complex bound to the polymerizing actin (19), and a debranching factor such as coronin in complex with F-actin, respectively. Mass conservation of + in the model could hence be attributed to competition for limited supply of actin or nucleating factors (59–64). In line with the global constraint, macropinocytic cup formation is known to compete with pseudopod formation (40). Appearance of an active patch on one side of the plasma membrane excludes another patch from appearing on other locations (65). As for the variable ***I***, its simulated profile (fig. S6*B*) is in line with that of coronin which trails behind the traveling actin waves (66). Apart from the role of ***I*** to inhibit amplification of ***A*** (Eq. 2), our model assumed ***I*** to increase by duplex of ***A*** (Eq. 3). In this regard, ***I*** can be regarded as a complex of coronin cross-linked with actin filaments (67, 68). In the activated patch, coronin may mediate the switch in the orientation of the dendritic actin filaments from those facing the membrane to those that are parallel (47). Such change would lead to vanishing force in the direction normal to the membrane interface consistent with our assumption that the force generation by ***A*** is competitively attenuated by ***I*** (Eq. (9)).

The current framework should be applicable to other related form of membrane deformation. Dendritic cells exhibit numerous multi-layered membrane ruffles and macropinosomes_(69, 70). Such coexistence of multiple internalized vesicles was rarely observed in the present simulation due to minimization of the global surface area assumed in the Allen-Cahn type phase-field equation (Eq. 1). Further exploration in the parameter space, specifically for large *a*_t_ and *α*, with perhaps additional implementation to control the speed of the vanishing vesicles may uncover related morphological features. In some cancer cells, the dorsal side of the plasma membrane is covered by circular membrane ruffle associated with macropinocytosis (19, 22). This so-called “circular dorsal ruffles” (CDR) is initiated from a F-actin-rich circular projections on the dorsal cell surface. Similar to the present simulations, the ring region expands then contracts, then forms a cup-like structure. Restriction of the dynamics in the dorsal side can be explained in the presence of dorsal-ventral asymmetry in the parameter at Phase I (e.g., Fig. 2*A*). We should note, however, that because multiple macropinosomes can form within a single cup (19), there likely is an additional mechanism at play to form these smaller ruffles. Further extension of the model such as to incorporate local change in tension *η,* which likely depends on localized myosin I (71, 72) may help explain these dynamics. Spatial restriction of the patch-driven dynamics may also help explain ruffling with a linear geometry known in macrophages where membrane ruffles many near the cell edge fold back on itself to close the cup (6, 73). Spatially much finer filopodial projections that resemble a tent-pole are also known (8). Future work should address the relation between these distinct subcellular morphologies and the basic cup dynamics uncovered in this work.

## Methods

### Numerical simulations

Time evolution of equation for *ϕ*, ***A*** and ***I*** was numerically solved using the standard explicit Euler method with mesh size *dx* = 0.1 μm and *dt* = 4.0 × 10^−4^ sec. For ***A*** and ***I***, instead of solving Eqs.(4) and (5) directly, we computed the following equations

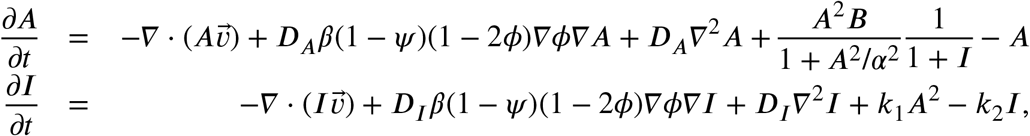

which derives from the relation 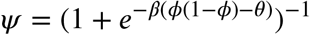. The above equations were solved on all lattice sites above the cut-off threshold *ψ* >10^−3^, otherwise ***A*** and ***I*** were allowed to simply decay at a rate *γ*_2_ = 10.0 [sec ^−1^]. Likewise, the equation for 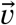 in Eq.(6) is computed for all sites |∇*ϕ*| > 10^−3^, otherwise 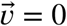. Note that, immediately after the cup closure, the internalized cup shrinks and eventually vanishes due to the surface tension which causes numerical instability due to an abrupt increase in ***A*** on the shrinking membrane. To avoid this instability, an upper limit was set to 50.0 for both ***A*** and ***I***. All simulations were coded in C. Results of three-dimensional simulations were visualized using OpenGL.

### Volume evaluation of the enclosed extracellular space

To reduce computation time, we considered a cell shape with z-axis symmetry so that the simulations can be run in the quasi 3-dimensional space with the z axis-symmetric coordinate (i.e., on a *z-r* plane). In this coordinate, ∇^2^ and 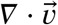 were replaced by 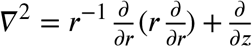 and 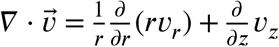. The Neumann boundary condition *∂_r_ϕ* = *∂_r_A* = *∂_r_I* = 0 are applied at the boundary *r* = 0, whereas Dirichlet boundary condition *ϕ* = ***A*** = ***I*** = 0 were applied for boundaries at ***z*** = 0, ***L_z_*** and *r = L_r_*, where ***L_z_, L_r_*** are the axial length of the system. The analysis consisted of two parts (fig. S1*G, H*): (1) scoring of the membrane enclosing events (i.e., whether or not the region with *ϕ* = 0 that is enclosed by *ϕ* =1 exists), and (2) estimating the enclosed volume at the time of cup closure. For the first part, for each simulation time step, the number of transition from *ϕ* = 0 to *ϕ* =1 (red circles in fig. S1*G*) was counted along the line ***r*** = Δ*r* from (Δ*r*, Lz) to (Δ*r*, 0). By definition, an enclosed region is present when this number is 4 (fig. S1*G*, right panel) otherwise no closure (fig. S1*G*, left panel). The enclosed volume was estimated at the time of closure by integrating the cross-sectional disk (fig. S1*H*, left panel) or disk with a hole at the center (fig. S1*H*, right panel) at constant z within *z_b_* ≤ *z* ≤ *z_t_*, where *z_t_* and *z_b_* are the first and second point at which *ϕ* changed from *ϕ* = 0 to 1 (fig. S1*H*).

## Acknowledgments

The authors thank Shuji Ishihara, Tetsuya Hiraiwa and Chikara Furusawa for helpful discussions. This work was supported by Japan Society for Promotion of Science (JSPS) Grant-in-Aid for Young Scientists JP18K13514 to NS, Japan Science and Technology Agency (JST) CREST JPMJCR1923, MEXT KAKENHI JP19H05801 to SS and in part by Joint Research by Exploratory Research Center on Life and Living Systems (ExCELLS) Grant 18-204, MEXT KAKENHI JP19H05416, JP18H04759 and JP16H01442; JSPS KAKENHI JP17H01812 and JP15KT0076 (to S.S.).

## Author Contributions

NS and SS conceived the work. NS planned the project, formulated the model, wrote and run the programs and performed all analysis. SS oversaw the project, supervised the analysis and contributed to the interpretation of the results. NS and SS wrote the manuscript.

