## Supplementary Information for "Three-dimensional morphodynamics simulations of macropinocytic cups"

**This PDF file includes:**

Supplementary text

Figures S1 to S8

Legends for Movies S1 to S5

**Other supplementary materials:**

Movies S1 to S5

**Supplementary Information Text**

**Phase field Implementation**

To simulate deformation of the plasma membrane a non-physical field $\phi$ is introduced. $\phi$ takes constant value ($\phi = 1$) at the cell interior region and ($\phi = 0$) at the exterior region, and varies sharply but smoothly in between. The width of the interface is characterized by the small parameter $\epsilon$, which can be interpreted as the thickness of the membrane and the cortical layer. The time evolution of the interface between $\phi = 0$ and 1 is considered by the general interface advection equation

$$\frac{\partial \phi}{\partial t} + \vec{v} \cdot \nabla \phi = 0,$$

where $\vec{v}$ is the velocity vector of the interface. We assume that the magnitude of the velocity is directly proportional to the force applied to the interface so that

$$\vec{v} = \frac{\vec{F} + \overrightarrow{F_c}}{\tau},$$

where the coefficient $\tau$ has the unit of [N$\cdot$ s/m$^3$], $\overrightarrow{F_c}$ is curvature-driven force normal to the plasma membrane that results from surface tension $\gamma$. Given the curvature $c$, $\overrightarrow{F_c} = -\gamma c \vec{n}$. In the phase field, the unit vector normal to the interface and curvature are given by $\vec{n} = -\nabla\phi/|\nabla\phi|$ and $-\nabla \cdot \vec{n} = \nabla \cdot (\nabla\phi/|\nabla\phi|)$. Following Beckermann et al (29), we employ a kernel function normal to the interface

$$\phi = \frac{1 - \tanh(\frac{\alpha n}{\epsilon})}{2},$$

where $n$ is the coordinate normal to the interface so that

$$c = \nabla \cdot \frac{\nabla\phi}{|\nabla\phi|} = \frac{1}{|\nabla\phi|}\left(\nabla^2\phi - \frac{\nabla\phi \cdot \nabla|\nabla\phi|}{|\nabla\phi|}\right)$$
$$= \frac{1}{|\nabla\phi|}\left[\nabla^2\phi - 4\frac{\alpha^2}{\epsilon^2}\phi(1-\phi)(1-2\phi)\right]$$

Thus we arrive at,

$$\tau\frac{\partial \phi}{\partial t} = \gamma(\nabla^2\phi - \frac{G'(\phi)}{\epsilon^2}) - \vec{F} \cdot \nabla\phi,$$

where $G = 2\alpha^2\phi^2(1-\phi)^2$ and $\vec{F} = M_V(\int \phi d\vec{r} - V_0)\nabla\phi/|\nabla\phi| - F_{poly}\nabla\phi/|\nabla\phi|$. The first force term represents the constraints for fixed cell volume and the second term is the force normal to the interface exerted by actin polymerization. Hence we obtain

$$\tau\frac{\partial\phi}{\partial t} = \gamma\left(\nabla^2\phi - \frac{G^{'}(\phi)}{\epsilon^2}\right) - M_V(V-V_0)|\nabla\phi| + F_{\mathrm{poly}}(\vec{r})|\nabla\phi|,$$

$$\vec{v} = -\left[\frac{\gamma\left(\nabla^2\phi - \frac{G^{'}(\phi)}{\epsilon^2}\right)}{|\nabla\phi|} - M_V(V-V_0) + F_{\mathrm{poly}}(\vec{r})\right]\frac{\nabla\phi}{|\nabla\phi|}.$$

**Kinetic equations for the active patch dynamics**

Eqs.(2) and (3) are derived as follows. For reaction $B \rightarrow A$ , we assume an autocatalytic reaction following a sigmoidal function with Hill coefficient 2 that is repressed by inhibitor $I$. For $A \rightarrow B$ , we assume a constant rate. The inhibitor is produced at the rate $k_1A^2$ and degraded at a constant rate $k_2$. Thus we write,

$$\begin{aligned}\frac{dA}{dt} &= k_A \frac{A^2B}{K_A^2 + A^2}\frac{1}{K_I + I} - d_AA + D_A\nabla^2A\\ \frac{dB}{dt} &= -k_A \frac{A^2B}{K_A^2 + A^2}\frac{1}{K_I + I} + d_AA + D_B\nabla^2B\\ \frac{dI}{dt} &= k_1A^2 - k_2I + D_I\nabla^2I,\end{aligned}$$

(S1)

which satisfies the conservation relation

$$\langle A\rangle + \langle B\rangle = a_{\mathrm{t}},$$

where $\langle\rangle$ indicates spatial average. By converting the variables to $\tilde{A} = \alpha A/K_A$, $\tilde{B} = \alpha B/K_A$, $\tilde{I} = I/K_I$, $\tilde{t} = d_At$ using a dimensionless parameter $\alpha = \sqrt{k_A/K_Id_A}$, we arrive at

$$
\begin{aligned}
\frac{d\tilde{A}}{d\tilde{t}} &= \frac{\tilde{A}^2\tilde{B}}{1+\tilde{A}^2/\alpha^2}\frac{1}{1+\tilde{I}} - \tilde{A} + \widetilde{D}_A\nabla^2\tilde{A} \\
\frac{d\tilde{B}}{d\tilde{t}} &= -\frac{\tilde{A}^2\tilde{B}}{1+\tilde{A}^2/\alpha^2}\frac{1}{1+\tilde{I}} + \tilde{A} + \widetilde{D}_B\nabla^2\tilde{B}, \\
\frac{d\tilde{I}}{d\tilde{t}} &= \tilde{k}_1\tilde{A}^2 - \tilde{k}_2\tilde{I} + \widetilde{D}_I\nabla^2\tilde{I},
\end{aligned}
$$

where $\widetilde{D}_A = D_A/d_A$, $\widetilde{D}_B = D_B/d_A$, $\widetilde{D}_I = D_I/d_A$, $\tilde{k}_1 = k_1K_A^2/k_A$, $\tilde{k}_2 = k_2/d_A$. Note that the spatial variable $\vec{r}$ $[\mu m]$ and the diffusion coefficients $[\mu m^2]$ still has dimension. Furthermore we reduce the number of kinetic variables to two by assuming that diffusion of $\tilde{B}$ is sufficiently fast in the timescale of our interest so that $\tilde{B}$ can be approximated as spatially homogeneous. Thus $\tilde{B}$ now becomes enslaved to $\tilde{A}$ according to the conservation relation

$$\langle\tilde{A}\rangle + \tilde{B} = \tilde{a}_{\mathrm{t}},$$

where $\tilde{a}_{\mathrm{t}} = a_{\mathrm{t}}\alpha/K_A$. Eqs (2) and (3) are obtained after redefining the variables and the parameters. Note that changes in the parameter in this reduced equation can interpreted by more than one of the original parameters. For example, when the reaction rate in $A \to B$ is lowered (i.e., decrease in $d_A$), due to mutation in Ras-GAP, $\alpha = \sqrt{k_A/K_I d_A}$ and $\tilde{a}_{\mathrm{t}} = a_{\mathrm{t}}\alpha/K_A$ increase, which results in enhanced active membrane patch and thus enhanced macropinocytosis and actin wave.

**Condition for the patch annihilation**

In the presence of the inhibitor, the activated spot with high $A$ can annihilates even without coupling with the membrane deformation. Whether the spot in Eq. (2) and (3) in the main text annihilates or not depends on the existence of a non-zero stable fixed point in the following equation:

$$
\begin{aligned}
\dot{a} &= \frac{a^2b}{1+a^2/\alpha^2}\frac{1}{1+I} - a \\
\dot{I} &= k_1a^2 - k_2I.
\end{aligned}
$$

At the fixed point, $a$ satisfies

$$0 = \frac{a^2b}{1+a^2/\alpha^2}\frac{1}{1+\kappa a^2} - a,$$

where $\kappa = k_1/k_2$. If and only if the function $f(a) = ab - (1 + a^2/\alpha^2)(1 + \kappa a^2)$ has a solution $f(a) = 0$ for $a > 0$ and $0 \le b \le a_t$, the non-zero fixed point exist in Eqs. (2) and (3). From $f(0) < 0$ and monotonicity of $f'(a)$, the condition for the absence of the non-zero fixed point for $0 \le b \le a_t$ is given by $f(a^*) < 0$, where $a^*$ satisfies $f'(a^*) = 0$. Thus we obtain

$$-\frac{1}{2}(\frac{1}{\alpha^2} + \kappa)(a^*)^2 + \frac{3}{4}a^* b - 1 < 0.$$

(S2)

The sufficient condition for patch annihilation is

$$\kappa > \frac{9}{32}a_t^2 - \frac{1}{\alpha^2}.$$

Note that neither a non-vanish static spot nor excitable behavior appears if this condition is met. The necessary and sufficient condition for patch annihilation would be obtained by numerically solving $f'(a^*) = 0$.

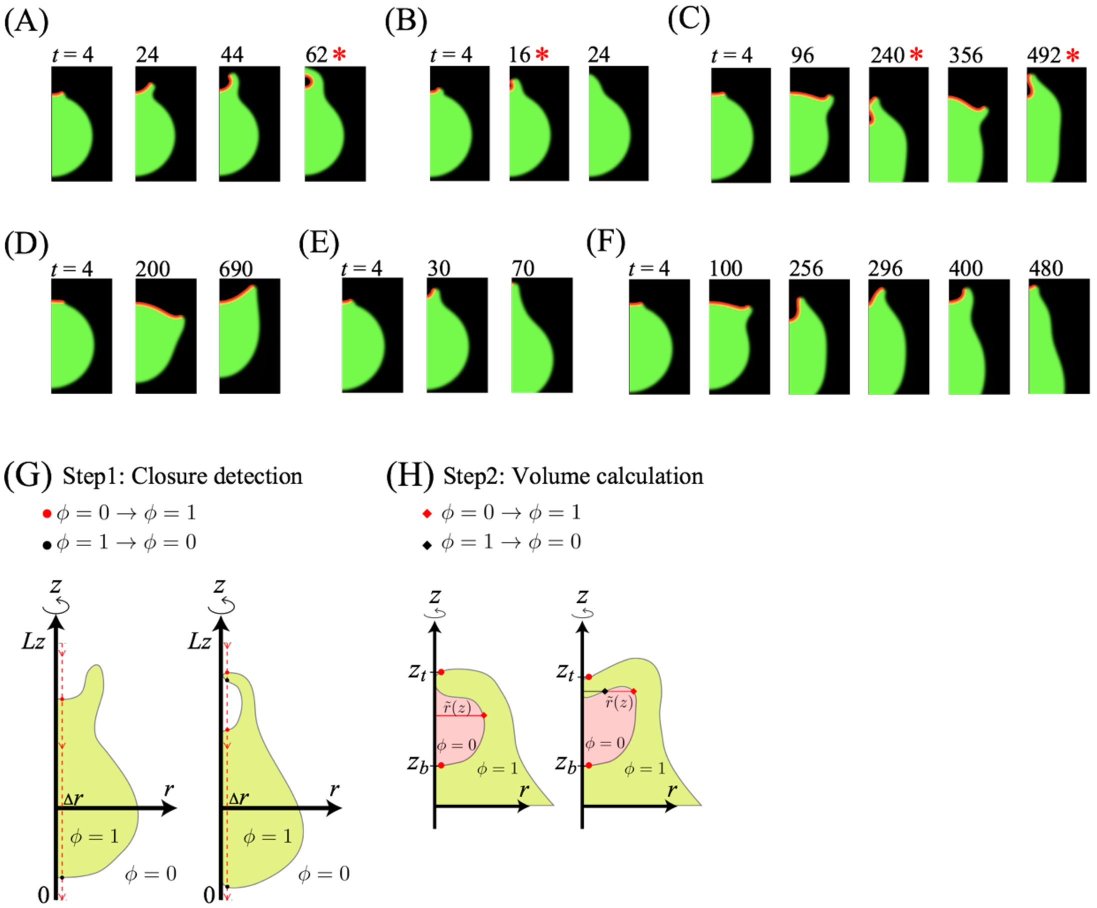


**Fig. S1: Calculation of the enclosed volume.** (*A-F*) Time course of active patch development in a quasi-3 dimensional space with z-axis symmetry. Parameters correspond to those used in (A) Fig. 2*A*, (B) Fig. 2*C*, (C) Fig. 2*F*, (D) Fig. 2*D*, (E) Fig. 2*E*, (F) Fig. 2*G*. (*G*, *H*) Calculation of the enclosed volume in the z-axis symmetric coordinate: (G) Detection of cup closure. The number of transitions between $\phi = 0$ and 1 (red and black circles) was counted along the linear path $r = \Delta r$ from ($\Delta r$, Lz) to ($\Delta r$, 0) where $\Delta r$ is the simulation mesh size (red dotted line). The first occurrence of 4 transitions was scored as the time of closure (right panel). (*H*) The enclosed volume was calculated by integrating the cross sectional area within $z_b \le z \le z_t$ (red shaded region).

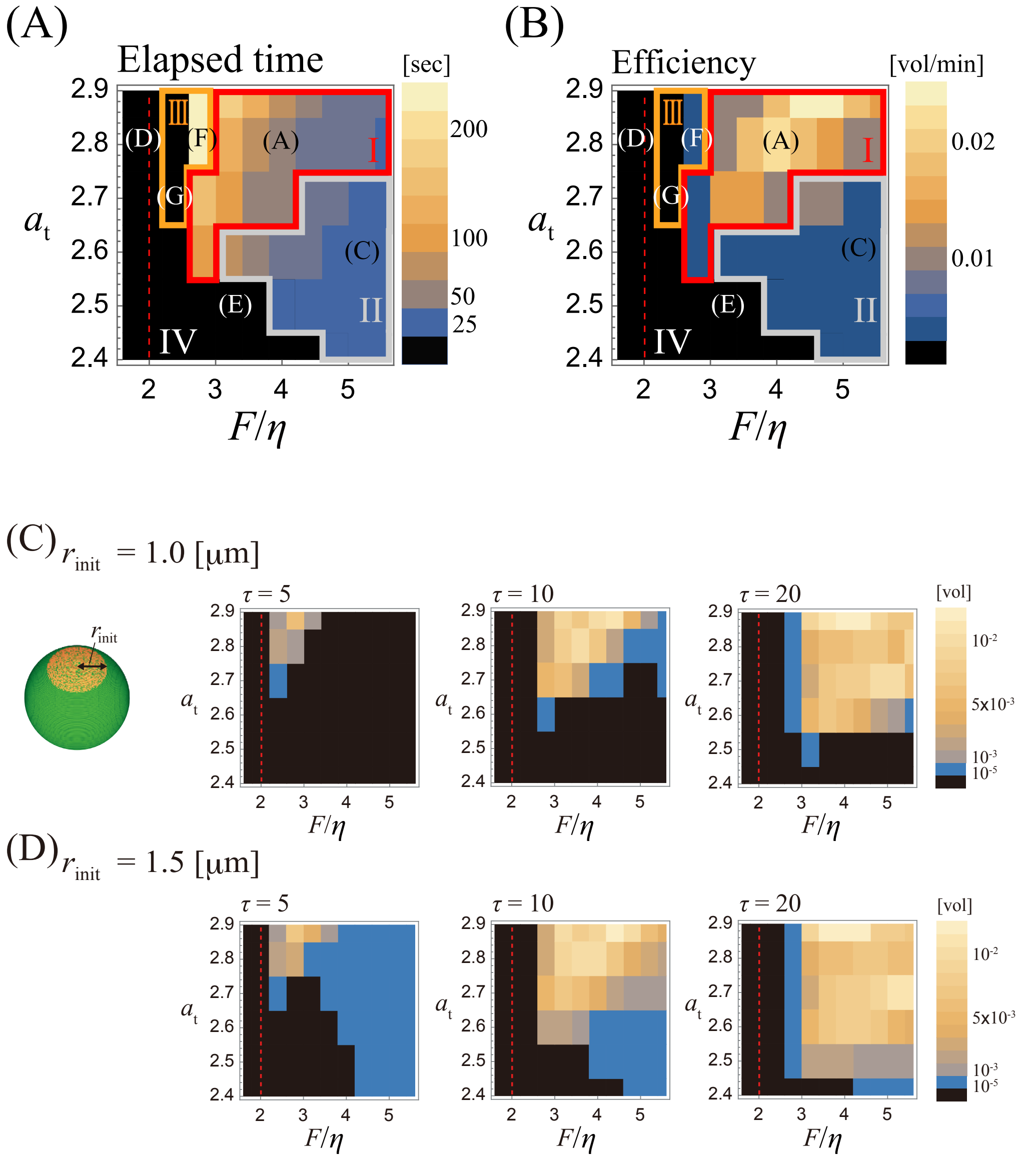


**Fig. S2: Cup formation and closure depends on the balance between the patch size and the strength of the protrusion force.** (*A*, *B*) Phase diagrams obtained from the quasi-3 dimensional space simulations. the elapsed time between patch initiation and cup closure (*A*) and the intake efficiency (*B*) defined by the normalized intake volume divided by the elapsed time. See Fig. 2*B* for the intake volume. Colors represent the averages of six independent simulation runs (three of each for $r_{\mathrm{init}} = 1.0$ μm and 1.5 μm) with successful cup closure. Parameter sets for Fig. 2*A* and C-G are indicated in the respective letters. (*C*, *D*) Phase diagram before averaging over $r_{\mathrm{init}}$. (*C*) $r_{\mathrm{init}} = 1.0$ and (*D*) $r_{\mathrm{init}} = 1.5$. Colors represent the average enclosed volume normalized by the total cell volume (blue to orange). Averages were obtained from three independent simulation runs. Enclosure of volume fraction smaller than $10^{-5}$ was scored as no uptake (black). Other parameters are the same as in Fig. 2.

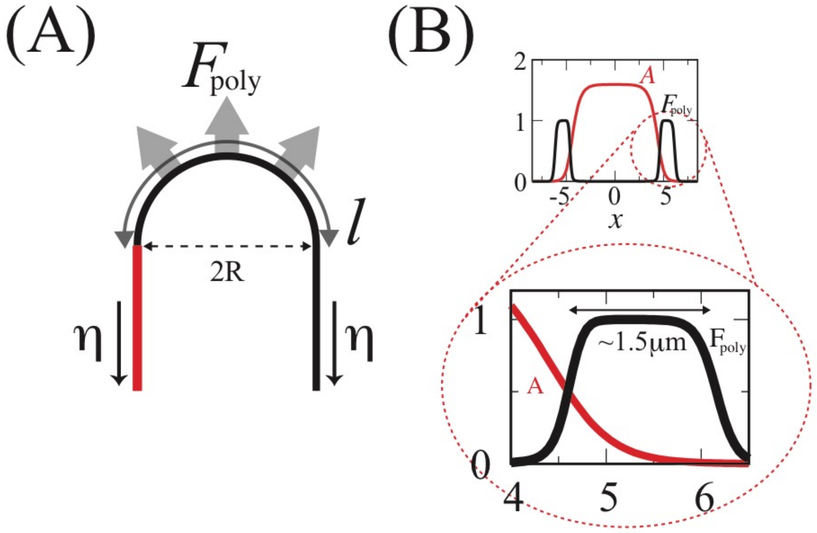


**Fig. S3: Outward protrusion force required to sustain cup-shaped membrane protrusion can be estimated from an approximated edge geometry**. (*A*) Force profile along the mid-line cross section of an idealized protrusion with width *2R* that consists the rim of the cup . Outward active force per unit length $F_{\mathrm{poly}}$ and the line tension $\eta$ are exerted on the semicircular head of length $l = \pi R$. Red represents a region with high $A$. (*B*) A blow-up view of the force distribution in Fig.1*D*. A snapshot from quasi-2 dimensional space simulation of Eqs. (2) and (3) ($a_t = 2.6$, $k_1 = k_2 = 0$, $\alpha = 1.0$, $D_a = 0.1$ and $D_i = 0.01$) without membrane deformation. Force in a direction normal to the membrane is exerted at the outer edge of width $l$ (a high $A$ region) ($F_{\mathrm{poly}}(A) \simeq F$ in Eq.(7)) . $l \sim 1.5$ [$\mu$ m] for $K_1 =$ 0.005, $K_2 =$ 0.25 and $n_{\mathrm{h}} =$ 3.

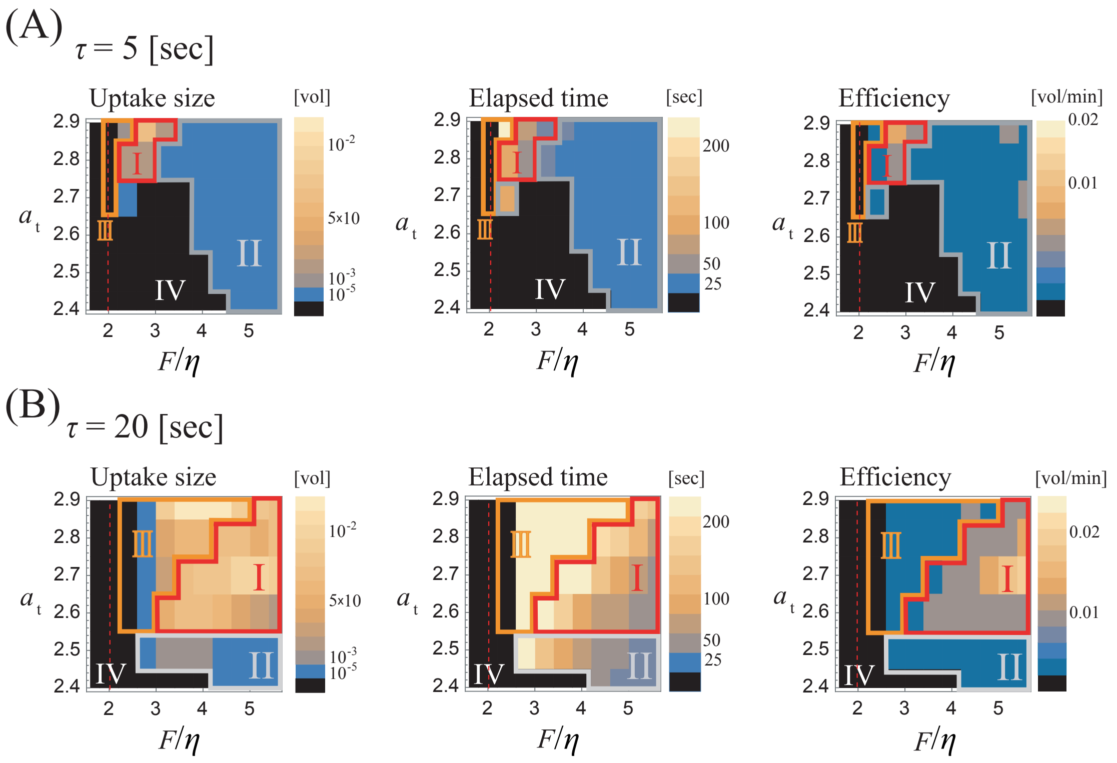


**Fig. S4 Deformation timescale acts critically for the intake volume and the elapsed time for cup closure.** (*A*, *B*) The average fraction of the enclosed volume (left panels), the elapsed time from the patch initiation to cup closure (middle panels), and the average fraction of the enclosed volume per minute (right panels). (*A*) $\tau = 5$ and (*B*) $\tau = 20$. The average of six independent simulations (three of each $r_{\text{init}} = 1.0$ μm and 1.5 μm). The enclosed volume smaller than $10^{-5}$ was scored as no uptake . For the middle and right panels, averages are taken from successful enclosure. For phase III, only the last of the repeated enclosure was sampled for averaging. Other parameters are the same as Fig.2.

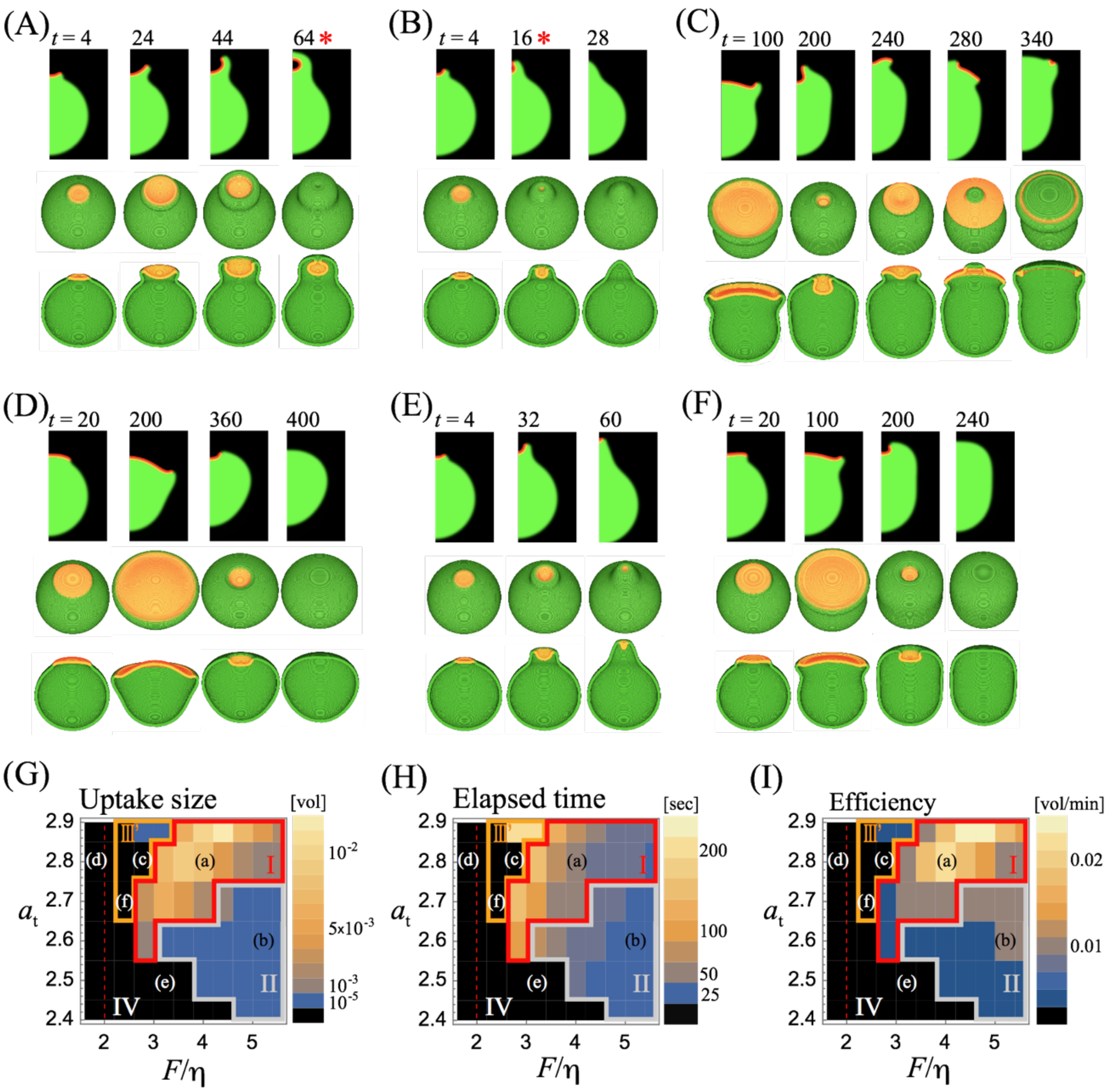


**Fig. S5: The inhibitor suppresses the repetitive cup formation at Phase III.** ($k_1 = k_2 = 2.0 \times 10^{-4}$) (*A-F*) Representative timeseries for $\tau = 10$. (*A*) $F/\eta = 4.0$, $a_t = 2.8$, (*B*) $F/\eta = 5.2$, $a_t = 2.6$, (*C*) $F/\eta = 2.8$, $a_t = 2.8$, (*D*) $F/\eta = 1.6$, $a_t = 2.8$, (*E*) $F/\eta = 3.2$, $a_t = 2.5$, (*F*) $F/\eta = 2.4$, $a_t = 2.7$. Other parameters are same as in Fig. 2*B*. Simulations in quasi-3 dimensional space (top panels). The top and the cross section view of the full-3 dimensional simulations (middle and bottom panels). Asterisks represents the time of cup closure. (*G-I*) Phase diagram of the cup dynamics. The enclosed volume normalized by the cell size (G), the elapsed time between the patch initiation and cup closure (*H*), and the efficiency (*I*). The average of six independent simulations (three of each $r_{\text{init}} = 1.0$ μm and 1.5 μm). The enclosed volume fraction smaller than $10^{-5}$ was scored as no uptake. For (*H*) and (*I*), the average was taken from samples showing successful cup closure.

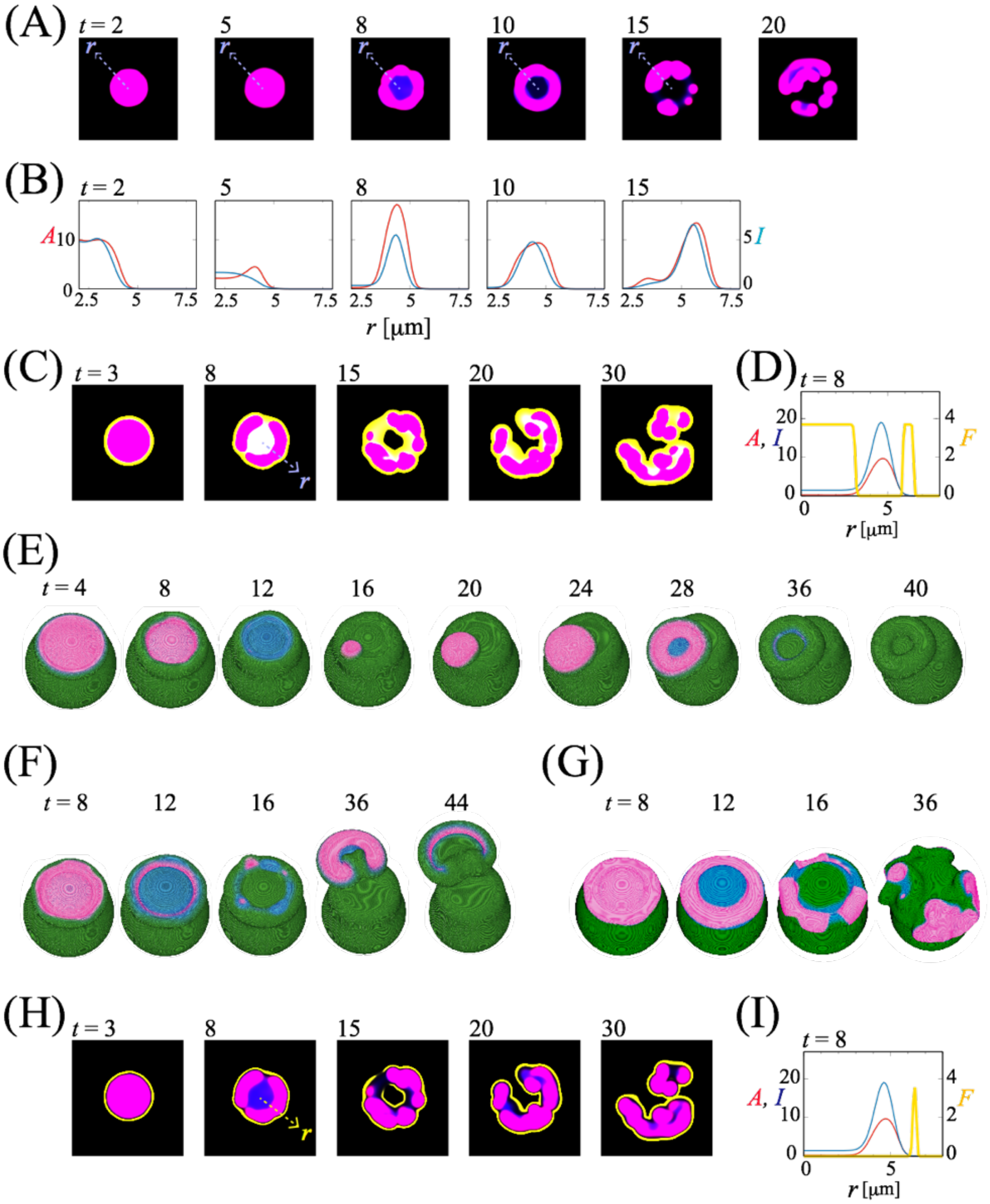


**Fig. S6: Splitting of patches and cups occur by destabilized wavefront in the presence of the inhibitor.** (*A-D*) Representative reaction dynamics on a flat 2-D space in the absence of membrane deformation for $a_t = 1.985$, $k_1 = 0.088$, $k_2 = 0.54$, $D_a = 0.085$, $D_i = 0.11$. Snapshots (*A*) and the cross-sectional profile (*B*) along the white dashed line in (A). (*C*, *D*) Imaginary force distribution computed according to $F_{\mathrm{poly}}$ with $A$ -dependency (Eq. (7)) (*C*; yellow). The spatial profile along the white line (*C*; $t = 8$)(*D*). (*E-G*) Representative membrane deformation simulations with Eq. (7); $F = 3.7$ (*E*), $F = 3.0$ (*F*) and $F = 2.0$ (*G*). Other parameters are same to Fig.4*C*. (*H*, *I*) Force distribution calculated according to protrusion force with $A$ and $I$ -dependency (Eq. (9)) (*H*; yellow).

Snapshots (*H*) and the cross-sectional profile (*I*) along the white line (*H*; $t = 8$). Colors in flat plane simulations indicate regions with overlapping ***A*** and ***I*** (pink) and ***I*** alone (blue), respectively (*A*, *C*, *H*). The color scheme for 3D simulations are the same as in Fig. 4.

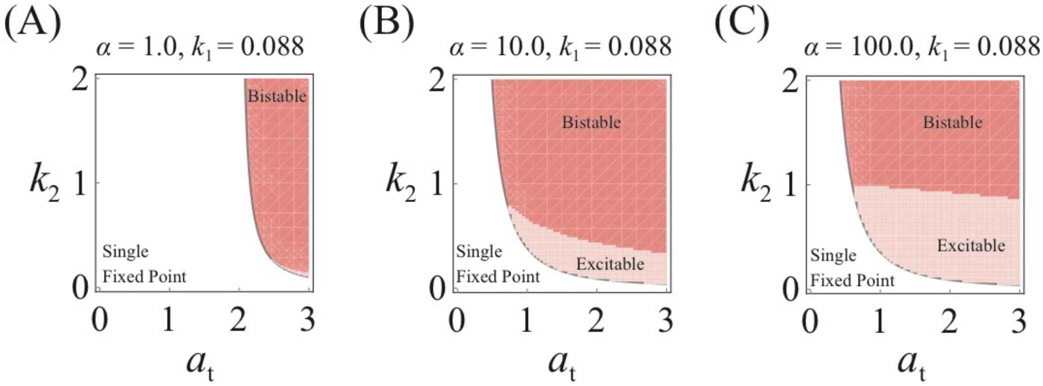


**Fig. S7: The excitable regime increases its domain in the parameter space for large $\boldsymbol{\alpha}$.** (*A*-*C*) Phase diagrams for Eqs. (2) and (3) at $k_1 = 0.088$ and $\alpha = 1.0$ (*A*), 10.0 (*B*) and 100.0 (*C*). Mono-stable (white): a patch vanishes and the spatially uniform state is stable at the steady state. Bi-stable (red): a spatial domain with high *A* coexist with low *A* domain at the steady state. Excitable (pink): small perturbation elicits large increase in A before returning to low *A* steady state.

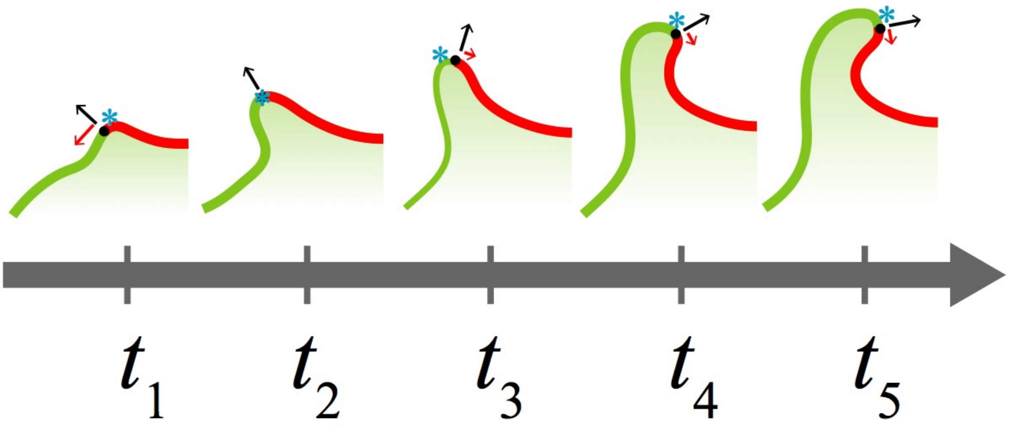


**Fig. S8 The edge of the active patch is displaced from the cup ridge due to limiting patch factor .** The protruding force is exerted at the edge (black circle) of a self-organized active patch (red). The protrusive force is normal to the membrane (black arrows). Expansion of an active patch ($t = t_1$; red arrow) slows down as the patch size grow and slightly exceeds its maximum size limit ($t = t_2$). As the patch begins to shrink and restores the maximum size, the edge of the patch (black circle) is slightly displaced towards the inner side of the cup, rather than at the very rim (highest curvature region, blue asterisk) ($t = t_3, t_4, t_5$; red arrows). When the protrusion wins over the surface tension that tries to restore high curvature in the inner cup region, the inward extension can grow to enclose a large volume of extracellular fluid until the edges meet and closes the cup.

**Legends for Movies S1 to S5**

Movie S1 (separate file). **Cup formation and closure from a single founder patch.** Timelapse images for the representative simulation result ($F/\eta = 4.0$, $a_t = 2.8$, $r_{\text{init}} = 1.5$ μm) shown in Fig. 2*A*. Merged RGB images : (red, $A\psi > 0$ (patch); green, $\psi > 0$ (membrane)).

Movie S2 (separate file). **Repetitive cup formation with cup closure at the waist.** Timelapse images for the representative simulation result ($F/\eta = 2.8$, $a_t = 2.8$, $r_{\text{init}} = 1.5$ μm) shown in Fig. 2*F*. Merged RGB images: (red, $A\psi > 0$ (patch); green, $\psi > 0$ (membrane)).

Movie S3 (separate file). **Repetitive cup formation with incomplete cup closure.** Timelapse images for the representative simulation result ($F/\eta = 2.4$, $a_t = 2.7$, $r_{\text{init}} = 1.5$ μm) shown in Fig. 2*G*. Merged RGB images (red, $A\psi > 0$ (patch); green, $\psi > 0$ (membrane)).

Movie S4 (separate file). **Multiple cup dynamics from stochastic patch initiation.** Timelapse images from a representative simulation with stochastic activation of $A$ ($F/\eta = 4.0$, $a_t = 2.8$, $\sigma = 8.0, d = 15.0, \lambda = 3 \times 10^{-5}$). Other parameters are same as in Fig. 2*A*. Merged RGB images (red, $A\psi > 0$ (patch); green, $\psi > 0$ (membrane)).

Movie S5 (separate file). **Cup splitting dynamics.** Timelapse images from a representative simulations with $I$ ($\tau = 7.0$, $F = 3.7, K_1 = 0.01, K_2 = 0.1$ and $n_h = 5$). Other parameters are the same as in Fig. 4. Merged RGB images (red, $A\psi > 0$ (patch); blue, $I\psi > 0$ (inhibitor); green, $\psi > 0$ (membrane)).
